# Silicon regulation of pectin methyl-esterification and cellulose microfibril organization modulates FERONIA-mediated cell wall integrity signaling for rice adaptation to salt stress

**DOI:** 10.1101/2025.03.18.644059

**Authors:** Yuyan Lei, Yuan Liu, Jing Wei, Wenbing Li, Shaoshan Zhang, Zhengming Yang, Jingqiu Feng, Ying Li, Huachun Sheng

## Abstract

The beneficial element silicon (Si) plays a crucial role in mitigating salt stress in plants and can activate intracellular signal transduction pathways that reprogram the transcriptome of salt-stressed plants. However, the mechanisms by which plants perceive Si nutrition, induce the expression of Si-responsive genes, and regulate the activity of Si-responsive proteins remain inadequately understood. In this study, we found that Si enhances cell wall integrity by increasing the degree of pectin methyl-esterification and changing the cellulose microfibril arrangement and functions as a regulatory switch for the activities of cell wall integrity sensors (FER homologs OsFLR1 and OsFLR2) during rice plant adaptation to high salinity. Furthermore, the inhibition of OsFLR1 and OsFLR2 through FER-specific inhibitors could negate the beneficial effects of Si and affect the uptake and accumulation of Si in rice seedlings. In conclusion, these findings suggest that Si signaling initiation involves FER-mediated cell wall integrity signaling pathways during salt stress adaptation in rice.

## 1. Introduction

The plant cell walls, composed of cellulose, hemicellulose, pectin, lignin, and structural proteins, provide a physical barrier that separates plant cells from their environment. Consequently, they protect plant cells against environmental stresses such as high salinity, and damage to cell wall integrity results in decreased salt stress tolerance [1, 2]. For instance, arabinose serves as a structural unit in several cell wall components and glycoproteins, and mutations in the UDP-arabinose synthesis-related gene *MUR4* render plants hypersensitive to salt stress due to an intrinsic mechanism whereby polysaccharides containing arabinose residues are essential for maintaining cell wall integrity under high-salinity conditions [3]. Additionally, defects in cellulose synthesis and excessive accumulation of β-1,4-galactan within the cell wall disrupt cell wall integrity and further diminish plant salt tolerance [4, 5].

Furthermore, cell wall integrity-sensing mechanisms have been proposed and several families of receptor kinases are identified as cell wall integrity sensors in plants, such as the *Catharanthus roseus* receptor-like kinase 1-like (CrRLK1L) family, wall-associated kinases (WAKs), and leucine-rich repeat receptor-like kinases (LRR-RLKs) [6, 7]. FERONIA (FER), a member of CrRLK1L family in Arabidopsis, is a well-known cell wall integrity sensor involved in plant tolerance to high salinity, which can interact directly with the stress-induced de-methyl-esterified pectic polysaccharides and then transduce the environmental cues to intracellular signals [8]. Very recently, it was proposed that FER mediated-phosphorylation of companion of cellulose synthase1 (CC1) controls microtubule array behavior in response to salt stress, by modulating CC1 trafficking and affecting the ability of CC1 to engage with microtubules [9]. There are two FER homologs with the highest homology in rice plants, OsFLR1 and OsFLR2, involved in root immunity and plant architecture [10, 11]. To date, no studies have shown that they are related to the rice response to salt stress.

All terrestrial plants contain a certain amount of silicon (Si), with most Si deposited in the epidermal cell walls and silica cells as hydrated SiO□ [12] while trace amounts of Si covalently crosslink with cell wall components as organosilicon [13, 14]. Si association and interaction with plant cell wall can alter the mechanical properties, surface potential and composition of plant cell wall [14-16]. Rice is a typical Si accumulator, with Si content reaching up to 10% of its dry weight, and it is also one of the most salt-sensitive cereals [17]. In salt-stressed rice plants, Si deposition into the root cell wall and Si-enhanced formation of Casparian strips can effectively block the apoplastic bypass flow of Na^+^, thereby reducing shoot Na^+^ concentration [18-20]. This suggests that Si alleviates ionic toxicity through an apoplastic obstruction mechanism. Additionally, Si nutrition stimulates the Salt-Overly Sensitive (SOS) pathway in the SOS1-dependent manner, further mitigating ionic toxicity in rice [21]. Beyond Na^+^ toxicity, salinity-induced osmotic stress can also be mitigated by Si regulation of root morphological traits and root osmotic potential, as well as Si promotion of transpiration force in the shoot [22]. Furthermore, Si increases the activity of anti-oxidative enzymes, which scavenge reactive oxygen species (ROS), leading to reduced oxidative damage and enhanced photosynthetic rates [22]. Kim et al. (2014) reported that prolonged exposure to increased Si concentrations can mitigate salinity-induced stress through the modulation of phytohormonal responses [23].

To sum up, Si plays a significant role in alleviating salt stress in plants by regulating ion and ROS homeostasis, enhancing root hydraulic conductance, and improving the metabolism of osmoregulatory substances and phytohormones, as well as increasing photosynthetic rates and the physicochemical properties of cell walls [24, 25]. Literature reviews indicate that the integration of Si with ROS metabolism is part of a signaling cascade that regulates phytohormone biosynthesis and morphological, biochemical, and molecular responses [25, 26]. However, how plants sense Si nutrition, induce the expression of Si-responsive genes, and regulate the activity of Si-responsive proteins remains poorly understood. Given the role of Si in modifying plant cell walls and the importance of cell wall integrity in salt tolerance, we hypothesize that Si signaling may initiate cell wall integrity signaling pathways during salt stress adaptation in rice. In this study, we provide evidence that Si regulates pectin methyl-esterification and cellulose microfibril organization to enhance cell wall integrity and influences FER-mediated cell wall integrity signaling to mitigate salt stress in rice plants.

## 2. Materials and Methods

### 2.1 Plant material and growth conditions

The japonica rice (*Oryza sativa* L.) variety Zhonghua 11 was utilized in this study. Rice seeds were initially treated with 75% ethanol for 2 minutes, followed by disinfection with 1% NaClO for 30 min, and finally immersed in deionized water for an additional 30 min. Subsequently, the seeds were soaked in deionized water and incubated in a constant temperature chamber at 30°C for two days to promote germination. Following seed germination, rice seedlings exhibiting consistent growth were selected and cultivated in Yoshida solution for 5 days before being transplanted into fresh nutrient solution with various treatments. The concentrations of Si (Na_2_SiO_3_·9H_2_O, Sigma-Aldrich, St. Louis, MO, USA), NaCl, and FER inhibitors (reversine and staurosporine; AbMole, Houston, TX, USA) were set at 1.5 mM, 150 mM, and 5 μM respectively. The rice seedlings were grown in an artificial climate chamber under the following conditions: a light/dark cycle of 16 hours/8 hours; relative humidity of 65%; day/night temperatures of 30°C/22°C; and light intensity of 450 μmol m□² s□¹.

### 2.2 Root cell wall chemistry analysis

For cell wall separation, 5-day-old rice seedlings were transplanted into fresh Yoshida solution containing Si, NaCl, or a combination of both. Four days later, the roots of the rice plants were harvested and ground to a powder in liquid nitrogen. The samples were vacuum-dried overnight and subsequently subjected to successive washes with 70% ethanol, a 1:1 (v/v) chloroform/methanol mixture, and acetone, each followed by centrifugation at 10,000 rpm for 10 minutes. After another round of vacuum drying, the cell wall extracts were prepared for subsequent analysis.

#### Fourier-transform infrared microscopy

The FT-IR spectra were obtained using a Thermo Nicolet iN10 FTIR spectrometer, utilizing cell wall extracts (100 mg) prepared by compressing a blend of fine dust with potassium bromide (KBr; 100 mg) powder. The wave number data derived from FTIR spectroscopy of cell walls, along with their corresponding assignments to the primary components of the cell wall, were presented in Table S1 [27].

#### Determination of the Content of Crystalline Cellulose

2 mg of cell wall powder were resuspended in 400 µL of a 2 M trifluoroacetic acid (TFA) solution and subjected to incubation at 121°C for a duration of 90 minutes. The material that remained insoluble in TFA was subsequently isolated, dried, and then redispersed in 1 mL of Updegraff reagent. After being incubated at 100°C for 30 minutes in boiling water, the samples underwent centrifugation at a force of 2500 g for 5 minutes. The residual crystalline cellulose was then dissolved in a solution containing 72% (v/v) H_2_SO_4_. Finally, the obtained crystalline cellulose was quantified colorimetrically using anthrone reagent at a wavelength of 625 nm with the aid of a spectrophotometer.

#### Pectin Content Measurements

To extract pectin, 2 mg of the cell wall extract was boiled with 1 mL of ultrapure water three times for 1 hour each. After each boiling, the supernatant was centrifuged at 16,800 g for 3 minutes and collected in a single tube. Galacturonic acid served as the standard to quantify the pectin content. A volume of 50 μL of the aforementioned pectin extract was added to 250 μL of 98% H_2_SO_4_ containing 0.0125 M Na_2_B_4_O_7_ and incubated at 100°C for 5 min in boiling water. Once cooled to room temperature, 5 μL of a solution containing 0.15% (w/v) meta-hydroxybiphenyl dissolved in 0.5% NaOH was added to the samples and incubated at room temperature for an additional 20 minutes. The absorbance of these samples at a wavelength of 520 nm was measured using a microplate reader.

#### Determination of the degree of pectin methyl-esterification

1 mg of cell wall extracts were initially washed with ultrapure water and centrifuged at 16,800 g for 10 minutes at room temperature to remove the supernatant. The samples were then incubated with 200 μL of NaOH at room temperature for 1 hour for pectin de-methyl-esterification, followed by adding 200 μL of 1 M HCl to stop the process. Subsequently, 125 μL of the reaction mixture was extracted and combined with 125 μL of a 20 mM HEPES solution (pH 7.5). Then, 250 μL of a HEPES solution containing 0.03 units of ethanol oxidase (Sigma-Aldrich) was added and incubated at room temperature for 15 minutes. After the reaction was done, an additional 250 μL of assay reagent (comprising 20 mM acetoacetone, 50 mM acetic acid, and 2 M ammonium acetate) was introduced and incubated at 60 °C for 15 minutes. After cooling to room temperature, absorbance at 412 nm was measured using a microplate reader. Methanol was used to construct a calibration curve to quantify methyl-esterification levels.

#### Hemicellulose Content Measurements

Hemicellulose extraction was performed on the residue obtained from pectin extraction. An aqueous solution containing 4% KOH and 0.02% KBH_4_ (1 mL) was added to the residue, and the resulting suspension was incubated at room temperature for 12 hours. Subsequently, the supernatant was collected by centrifugation at 16,800 g for 10 minutes at room temperature. This procedure was repeated once more, and the extracts from both rounds were combined to yield the hemicellulose extract. The hemicellulose content was quantified using glucose as a standard via the phenol-sulfuric acid method. To this end, 1 mL of 98% H_2_SO_4_ and 10 μL of an 80% phenol solution were added to 200 μL of the hemicellulose extract. The samples were initially incubated at room temperature for 15 minutes before being boiled for another 15 minutes. After cooling to room temperature, absorbance measurements of the samples were taken at a wavelength of 490 nm using a microplate reader.

### 2.3 Cell wall integrity assay

#### Propidium iodide (PI) staining

5-day-old rice seedlings cultivated on Yoshida medium with or without 1.5 mM Si were subsequently transferred to a medium enriched with 150 mM NaCl. Following a growth period of 16 hours, the roots were stained with PI prior to cell integrity evaluation using confocal microscopy (Zeiss, Oberkochen, Germany). Cell death was characterized by complete staining with PI. The number of burst cells in the elongation zone of the roots was quantified.

#### Atomic force microscopy

The protocol for atomic force microscopy (AFM) was adapted from the methodologies established by He et al. (2015) [28] and Pu et al. (2021) [29]. The cell wall powder was re-suspended in ultrapure water and subsequently deposited onto a poly-l-lysine-coated mica substrate. AFM imaging was conducted using the NanoScope 8 atomic force microscope (Bruker, Santa Barbara, CA) in ScanAsyst-Air mode. To assess mechanical properties, representative Force-Distance (F-D) curves were acquired and analyzed with Nanoscope analysis software version 1.9 (Bruker, Santa Barbara, CA). Young’s modulus was determined utilizing the Sneddon model as described by Sneddon (1965) [30].

### 2.4 RNA Isolation and Quantitative Real-Time PCR Analysis

5-day-old rice seedlings were transferred to new nutrient solutions containing various treatments. For RNA extraction, samples were collected at 12-hour and 24-hour intervals following the treatment. Total RNA was extracted using the RNeasy Plant Mini Kit (QIAGEN, Hilden, Germany). Complementary DNA (cDNA) was synthesized (Invitrogen, Carlsbad, CA, USA) and subsequently utilized as a template for quantitative real-time PCR (qRT-PCR), conducted on a CFX96 Real-Time System (Bio-Rad Laboratories, Hercules, CA, USA) in conjunction with the SYBR Premix Ex Taq II Kit (Takara Bio, Otsu, Japan). The rice *Histone H3* gene served as an internal control for normalizing the expression levels of target genes. The sequences of primers used for amplification in qRT-PCR analyses are provided in Table S2.

### 2.5 Si content measurements

100 mg of dried rice seedlings were ground into a fine powder, placed in a Teflon crucible, and combined with 6 mL of NaClO and 2 mL of 2 M NaOH. The crucible was subsequently heated at 150°C for 30 minutes to facilitate digestion of the sample. After cooling, the resulting solution was transferred to a 250 mL volumetric flask containing 9 mL of 1 M H□SO□ and 100 mL of distilled water, followed by dilution to the mark. A volume of 1.5 mL from this solution was then transferred to a separate 50 mL volumetric flask and mixed with an additional 2 mL of 98% H□SO□ until homogeneous. Five minutes later, another addition of distilled water (20 mL) was made, followed by thorough mixing with 5 mL of 5.0% ammonium molybdate solution and 2 mL of 0.5% ascorbic acid. The mixture stood for 20 minutes before being diluted to the mark, after which absorbance measurements were conducted at 810 nm.

### 2.6 Analysis of ABA content

Rice seedling was homogenized using a glass rod. Subsequently, the homogenates were each subjected to two rounds of methanol extraction, and the extracts were pooled separately. The pooled extracts were then centrifuged. The supernatants were passed through an HLB column, followed by passage through an MCX column. The collected samples were evaporated to dryness and re-dissolved in 500 μL of chromatographic-grade methanol for subsequent UPLC-MS/MS analysis (Shimadzu UPLC-MS/MS 8050, Kyoto, Japan). Moreover, a concentration gradient of ABA standard (5, 2, 1, 0.5, 0.2, 0.1, 0.05, 0.02, and 0.01 μg/mL) was prepared in chromatographic-grade methanol. MS detection was performed in multiple reaction monitoring (MRM) mode, and the compound-specific MRM parameters were set as previously reported [31].

### 2.7 Statistical analysis

For the measurements of plant height and fresh weight, all experiments were conducted at least in triplicate. The sample size, denoted as n, for each experiment is provided in the respective figure legends. For root cell wall chemistry and integrity analysis, qRT-PCR, and content measurements of Si and endogenous ABA, all experiments were performed at least three times using independent biological replicates, with consistent results observed across trials. All reported experimental data represent means ± standard deviation (SD) and were statistically analyzed using Welch’s one-way ANOVA.

## 3. Results

### 3.1 Si effects on pectin methyl-esterification and cellulose microfibril organization of root cell wall under salt stress condition

The effects of Si, NaCl, or their combination on cell wall chemistry were assessed using Fourier transform infrared (FTIR) spectroscopy and chemical methods. The highest peaks for cellulose (approximately 1050 cm^−1^) were observed in rice plants across all treatment conditions, indicating elevated cellulose levels (Fig. 1A). No significant differences were detected at the wavenumbers corresponding to hemicelluloses (between 1161 and 1442 cm^−1^), pectin (between 950 and 990 cm^−1^), and esterified uronic acid (1712 and 1735 cm^−1^) under the three treatment conditions (Fig. 1A). Specific measurements of hemicellulose, and pectin largely exhibited a similar trend as the FTIR results (Fig. 1C-D), and the cellulose content rose slightly but without statistical significance (Fig. 1B). Notably, Si appeared to mitigate the NaCl-induced de-methyl-esterification of pectin (Fig. 1E).

**Fig. 1.**
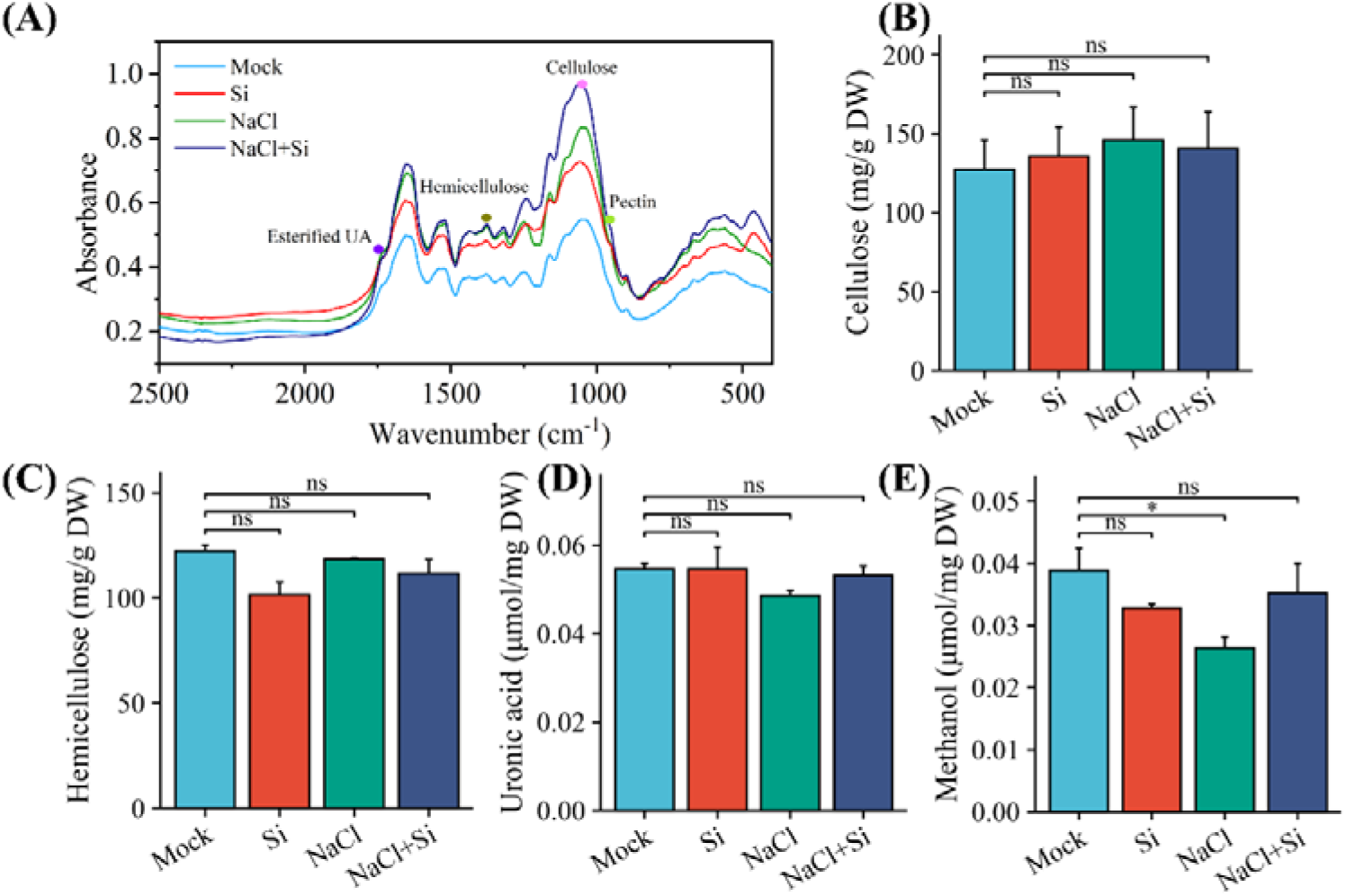
Si alters cell wall chemistry in a pectin-dependent manner in salt-stressed rice plants. (A) FTIR spectroscopy of root cell wall. (B-D) Cellulose, hemicellulose and uronic acid content measured using the colorimetric method. (E) Fractions of methylesterified pectic polysaccharides. Values are means ± SD (*n* = 3). Data were statistically evaluated using Welch’s one-way ANOVA, and the significance of differences (*, p < 0.05; ns, not significant) is indicated.

To further investigate the Si effects on cell wall integrity under salt stress conditions, NaCl-induced cell bursting and cell wall softening were detected by PI staining and AFM, respectively. It was observed that the addition of Si could reduce the NaCl-induced cell bursting (Fig. 2A, B). Moreover, the AFM results indicated that Si-modified cell walls possess denser cellulose microfibrils (Fig. 2C) and enhanced mechanical properties under high salinity conditions (Fig. 2D). According to the above, Si could maintain the cell wall integrity under salt stress conditions by increasing the methyl-esterification degree of pectin and changing the cellulose microfibril arrangement.

**Fig. 2.**
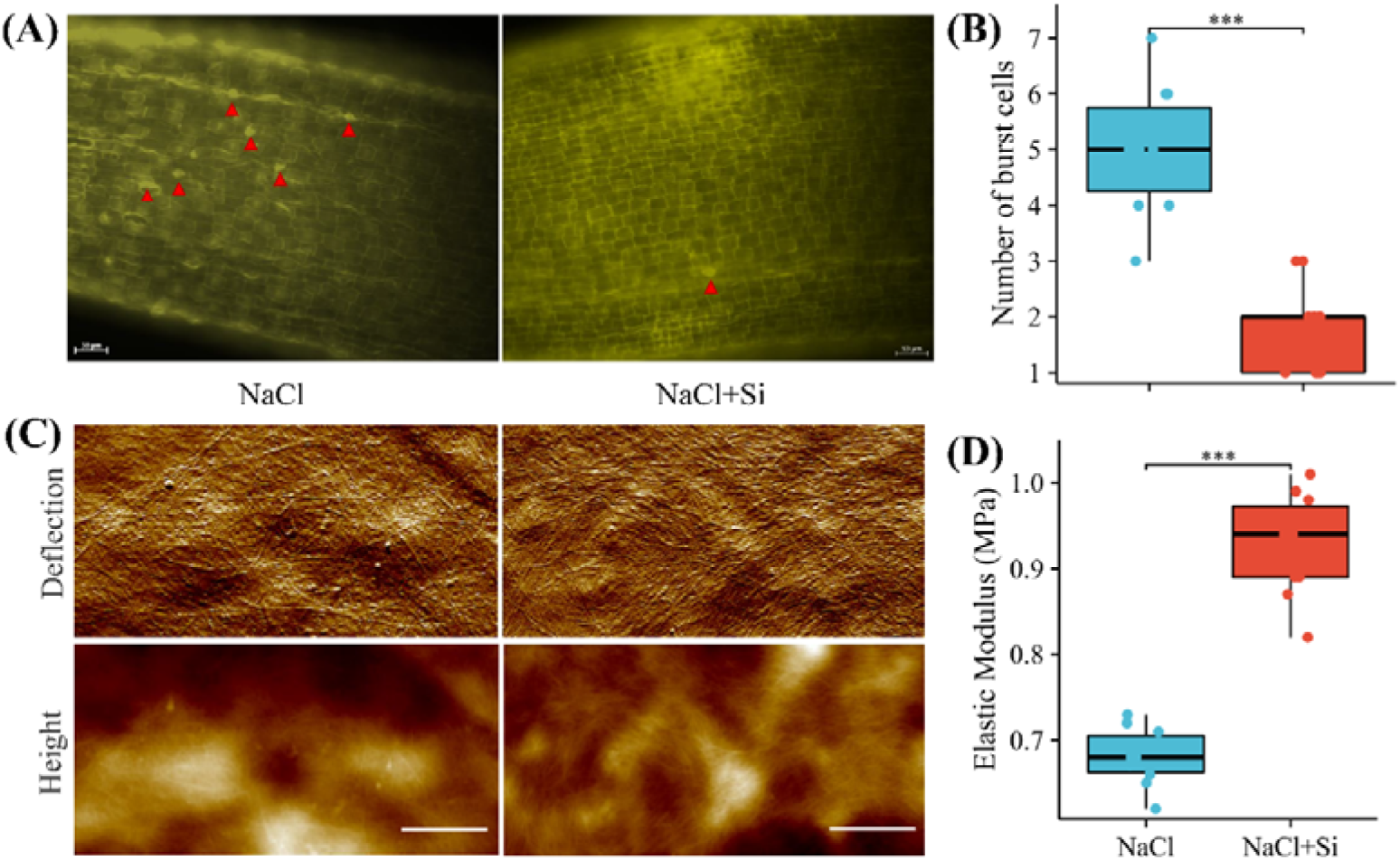
Si improvement of cell wall integrity. (A) Images of cells in the root elongation zone stained with PI. (B) The number of burst cells in the root elongation zone. (C) Images of root cell wall by atomic force microscopy. (D) Average Young’s modulus of the root cell wall. Values are means ± SD (n =10). Data were statistically evaluated using Welch’s one-way ANOVA, and the significance of differences (***, p < 0.001) is indicated. Scale bar = 50 μm (A) and 1 μm (C).

### 3.2 Si and NaCl influence the expression of FER homolog genes

In rice plants, OsFLR1 and OsFLR2 are the homologs of FER (Fig. 3A). Salt stress reduces the activity of FER kinase [32], while the expression of the FER gene is upregulated to sustain its activity in plant response to high salinity [33]. This also applies to its homologous genes *OsFLR1* and *OsFLR2* in rice (Fig. 3B, C), suggesting that the FER homologs OsFLR1 and OsFLR2 also play a role in salt stress responses. Notably, Si enhances the NaCl-induced expression of *OsFLR1* and *OsFLR2* genes during the early stages when rice plants are exposed to high salinity within 12 hours (defense phase), whereas after 24 hours of treatment (re-growth phase), Si suppresses the NaCl-induced expressions of these FER homolog genes (Fig. 3B, C). Considering that Si downregulates *OsFLR1* and *OsFLR2* gene expressions under normal conditions, it can be concluded that Si plays a dual role in regulating the expression of FER homolog genes, thereby mediating the trade-off between plant growth and defense.

**Fig. 3.**
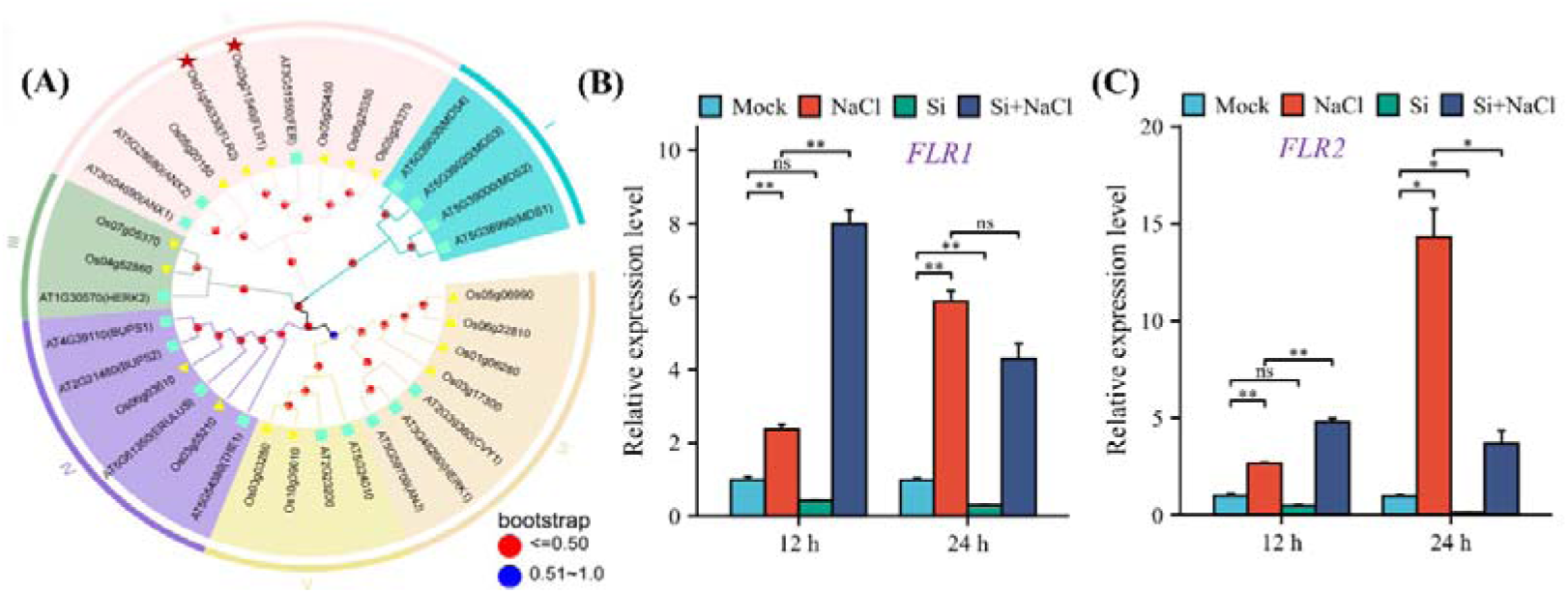
Relationship between Si and FER kinase activity. (A) Comparative phylogenetic analysis of rice and Arabidopsis CrRLK1L proteins. (B, C) The expression patterns of FER homolog gene *OsFLR1* and *OsFLR2* under the NaCl, Si, and Si+NaCl conditions. Values are means ± SD (*n* = 3). Data were statistically evaluated using Welch’s one-way ANOVA, and the significance of differences (**, p < 0.01; *, p < 0.05; ns, not significant) is indicated.

### 3.3 Si mitigation of salt stress depends on the activation of FER kinase

Under salt stress conditions, Si-mediated cell wall integrity maintenance may be perceived by FER homologs. To test this hypothesis, reversine and staurosporine were employed in this study as FER-specific inhibitors that can inactivate FER kinase and its homologs by targeting the conserved ATP binding pocket within the kinase structure [34]. The addition of reversine had no significant effects on plant growth (Fig. 4A-C), whereas staurosporine exhibited inhibitory effects on both plant height and fresh weight (Fig. 4G). Notably, Si was able to mitigate salt stress in rice seedlings; however, its beneficial effects were negated by the application of both FER inhibitors (Fig. 4D-J), suggesting that Si mitigation of salt stress is contingent upon the activation of FER kinase.

**Fig. 4.**
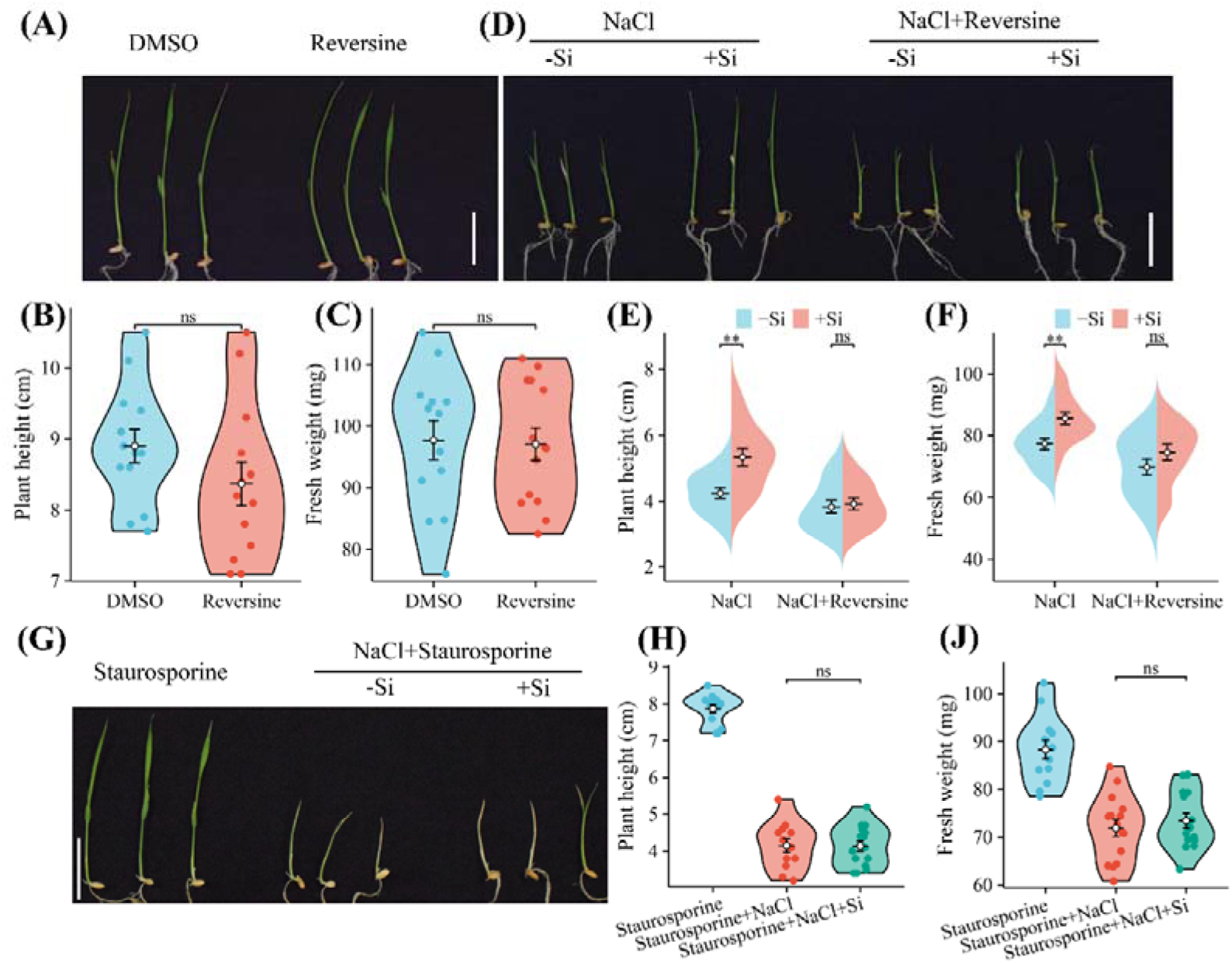
Si roles depend on the activation of FER kinase. (A) Images of hydroponic seedlings treated with reversine. 5-day-old rice seedings were transplanted into the medium containing reversine for 4 days. (B, C) Plant height and fresh weight of rice seedlings after being transferred to the medium with reversine for 4 days. (D) Images of hydroponic seedlings treated with NaCl-Si, NaCl+Si, NaCl-Si+reversine, and NaCl+Si+reversine. 5-day-old rice seedings were transplanted into the medium containing the aforementioned four treatments for 4 days. (E, F) Plant height and fresh weight of rice seedlings measured in those shown in (D). (G) Images of hydroponic seedlings treated with staurosporine, NaCl-Si+staurosporine, and NaCl+Si+staurosporine. 5-day-old rice seedings were transplanted into the medium with above three treatments for 4 days. (H, J) Plant height and fresh weight of rice seedlings measured in those shown in (G). Values are means ± SD (*n* > 12). Data were statistically evaluated using Welch’s one-way ANOVA, and the significance of differences (**, p < 0.01; ns, not significant) is indicated. Scale bar = 3 cm.

### 3.4 FER kinases could regulate the uptake of Si

To investigate the interactions between Si and FER, we assessed the expression of Si transporter genes (*OsLsi1* and *OsLsi2*) and Si uptake in rice seedlings treated with NaCl, a FER inhibitor (reversine), or both. The expression levels of *OsLsi1* and *OsLsi2* genes were downregulated by treatment with NaCl or the FER inhibitor, with a more pronounced effect observed when both treatments were applied simultaneously (Fig. 5A, B). We also measured Si content in rice seedlings subjected to NaCl, the FER inhibitor, or both and found that Si uptake was decreased under these conditions (Fig. 5C). Collectively, these findings suggest that FER homologs positively regulate the expression of Si transporter genes and the uptake of Si in rice plants.

**Fig. 5.**
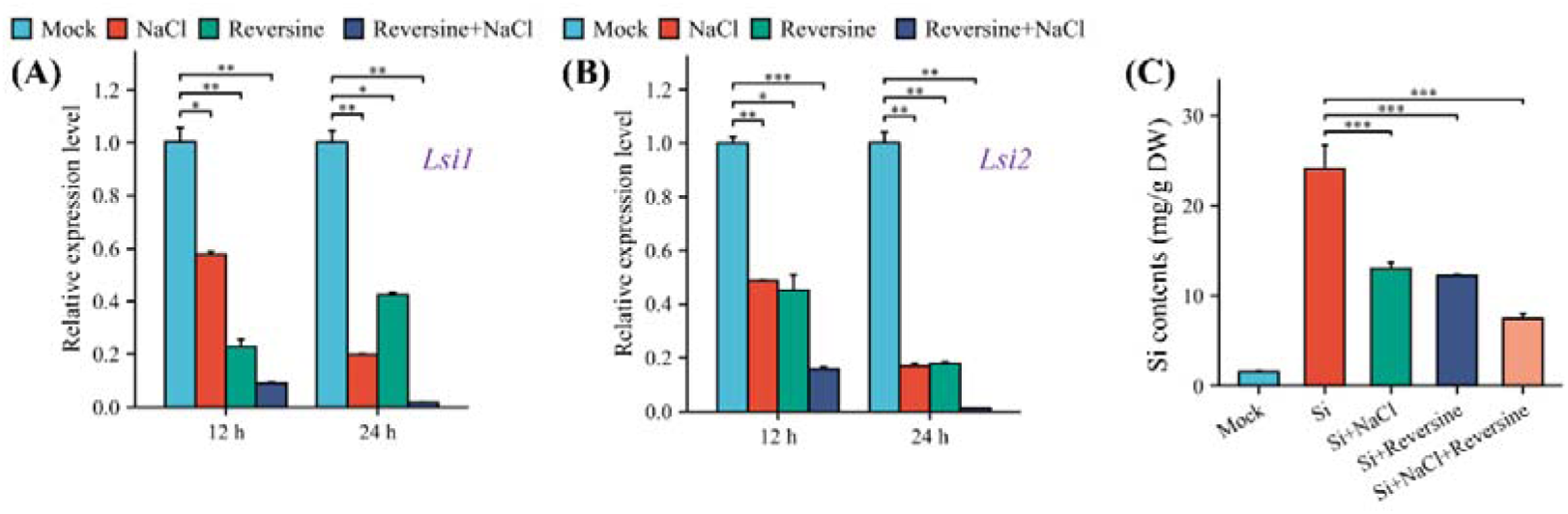
FER kinase regulates the uptake and accumulation of Si. (A, B) The expression patterns of Si transporter gene *OsLsi1* and *OsLsi2* under the NaCl, reversine, and Si+reversine conditions. (C) Si contents in rice seedlings treated with Si, Si+NaCl, Si+reversine, Si+NaCl+reversine. 5-day-old rice seedings were transplanted into the medium with above treatments for 4 days and then the Si contents were measured using the colorimetric method. Values are means ± SD (*n* = 3). Data were statistically evaluated using Welch’s one-way ANOVA, and the significance of differences (***, p < 0.001; **, p < 0.01; *, p < 0.05) is indicated.

### 3.5 ABA biosynthesis is mediated by Si and FER kinases

Previous studies have reported that the expression of *OsLsi1* and *OsLsi2* genes negatively correlates with endogenous abscisic acid (ABA) contents due to the presence of ABA-responsive motifs in their promoter regions [35]. Therefore, we detected the ABA contents under the normal and salt stress (Fig. 6) conditions. It has been reported that salt stress and FER inactivation lead to increased ABA production [33], and our results align with those previously reported. Interestingly, Si could markedly enhance ABA biosynthesis under the normal condition (Fig. 6A) but inhibit it under the long-term salt stress condition. Moreover, when FER is rendered inactive by its specific inhibitor, the role of Si is conspicuously attenuated under the long-term salt stress condition (Fig. 6B), suggesting Si regulation of ABA production in a FER activation-dependent manner. And, these findings are in accordance with the results shown in Fig. 4.

**Fig. 6.**
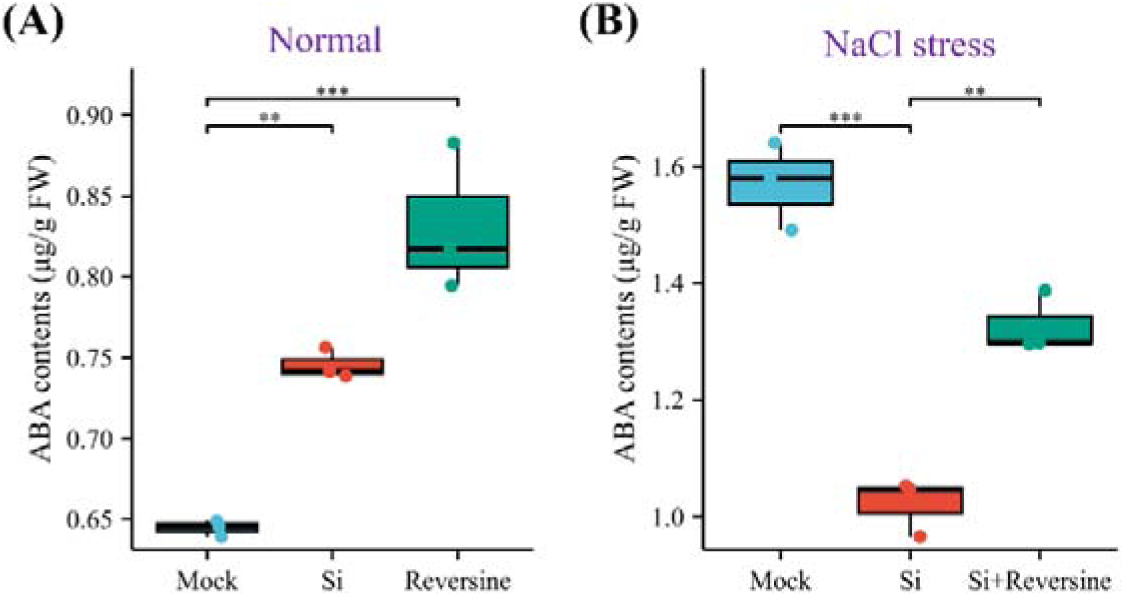
ABA biosynthesis is affected by Si and FER kinase under normal (A) and salt stress (B) conditions. 5-day-old rice seedlings were transplanted into fresh Yoshida solution containing Si, Reversine, NaCl, NaCl+Si, NaCl+Si+Reversine. Four days later, rice plants were harvested to detect ABA content. Values are means ± SD (n = 3). Data were statistically evaluated using Welch’s one-way ANOVA, and the significance of differences (***, p < 0.001; **, p < 0.01) is indicated.

## 4. Discussion

### 4.1 Si regulates pectin methyl-esterification and cellulose microfibril organization for cell wall integrity maintenance under salt stress conditions

It is well established that the cell wall plays a critical role in plant defense against environmental stresses including salt stress, and remodeling of the cell wall through alterations in its composition and organization constitutes a vital adaptive strategy [1, 2, 36]. Prior to the recovery of growth rates in salt-stressed plants, the cell wall particularly that of root epidermal cells, undergoes a transition from softening to stiffening [8]. Initially, excess sodium ions (Na^+^) can disrupt the Ca²□-crosslinked pectin network by competing with Ca²□ for negative charges on galacturonic acid residues [37], leading to a softened cell wall. To maintain cell wall integrity, pectic polysaccharides are subsequently de-methyl-esterified through the activation of pectin methylesterases (PMEs) in plants, which polysaccharides can then be cross-linked by divalent cations to restore cell wall stiffness [38, 39].

The beneficial element Si plays a positive role in alleviating salt stress in plants [24, 25], and its association and interaction with cell wall components have been proposed [13, 14]. Previous studies have regarded deposited silica as supplementary material that strengthens plant cell walls [40]. Si also collaborates with lignin to enhance cell wall stiffness: it can induce the biosynthesis of lignin, a component of the secondary cell wall; conversely, lignin interacts with mono-silicic acid to regulate silica deposition within the cell walls [41, 42]. Increased deposition of both lignin and silica forms apoplastic obstruction that prevent Na□ from entering cells [40]. In addition to inorganic silica, our research group focuses more on cell wall-bound Si. Recently, Pu et al. (2021) reported that xyloglucan serves as the ligand for bonding Si in rice cell walls, participating in establishing and remodeling the cell wall architecture through cross-linking with cellulose microfibrils, thereby contributing to increased stiffness [29]. It has been demonstrated that Si-xyloglucan complexes not only enhance resistance to enzymatic degradation and salt-alkali digestion but also improve the nanomechanical properties of rice plant cell walls [13, 29].

In addition to lignin and the hemicellulose xyloglucan, pectic polysaccharides can serve as silicification templates for SiO□ deposition [43] and crosslink with mono-silicic acid to form Si-pectin complexes [44]. Here, we aim to investigate the relationship between salt stress adaptation and Si-pectin interactions in rice plants. Our data indicate that Si application can counteract NaCl-induced de-methyl-esterification of pectin (Fig. 1), thereby rearranging cellulose microfibril organization and increasing root cell wall stiffness (Fig. 2) and suggesting that Si maintains cell wall integrity in the cellulose- and pectin-dependent manner. Similar results were observed in aluminum-stressed rice and lanthanum-stressed *Dicranopteris linearis*: the addition of Si increases the degree of pectin methyl-esterification and preserves cell wall integrity under stress conditions, thus facilitating recovery of plant growth [45, 46].

### 4.2 The initiation of Si signaling involves FER-mediated cell wall integrity signaling pathways in rice plants response to salt stress

Changes in cell wall composition and organization trigger the cell wall integrity signaling pathway [6, 7]. FER is a plasma membrane-localized receptor kinase with an extracellular ligand-binding domain that senses cell wall integrity, perceiving salt stress-mediated cell wall damage through binding to de-methylesterified homogalacturonan [8, 47]. In this study, we found that the FER homolog genes of *OsFLR1* and *OsFLR2* are induced by NaCl treatments (Fig. 3) and OsFLR1 and OsFLR2 positively regulate the salt stress tolerance in rice seedlings (Fig. 4). Moreover, recent reports indicate that the pectin-FER complex plays a role in plant mechanical force perception and activates ROP6 GTPase signaling during Arabidopsis pavement cell morphogenesis, suggesting that FER kinase also functions as a mechanoreceptor [48].

Considering the effects of Si on pectin methyl-esterification and the mechanical properties of rice plant cell walls, it appears that Si may influence FER-mediated cell wall integrity signaling under stress conditions [49]. Indeed, when the activity of FER kinase is inhibited by specific inhibitors, the beneficial effects of Si are abolished (Fig. 4), implying a direct link between Si and FER. Interestingly, Si acts as a regulatory switch for the activities of FER homologs OsFLR1 and OsFLR2 (Fig. 3): Si can enhance their activities during stress response stages and reduce them during growth recovery periods. Herein we propose a hypothesis that Si signaling initiation involves FER-mediated cell wall integrity signaling pathways during salt stress adaptation in rice plants.

Very recently, a homolog of the flowering hormone “florigen” FT, specifically FLOWERING LOCUS T-LIKE 12 (OsFTL12), has been identified as the Shoot-Silicon-Signal protein [50]. This protein is predominantly expressed in the shoot; however, under Si-deficient conditions, it also reaches the phloem and roots, thereby inducing increased expression of transporters (OsLsi1 and OsLsi2) responsible for Si uptake in roots [50]. It can be concluded that OsFTL12 is a potential component of the Si signaling process but not the receptor of Si signaling, because how low Si induces the expression of *OsFTL12* gene is still unknown. It is worth noting that FER modulates the flowering time by regulating the expression of *FT* in Arabidopsis and the lower peak in *FT* expression was observed in the *fer-4* mutant [51]. By coincidence, the inactivation of FER homologs OsFLR1 and OsFLR2 down-regulates the expression levels of Si transporter genes *OsLsi1* and *OsLsi2*, leading to decreased uptake and accumulation of Si in rice seedlings (Fig. 5). Similar results were also observed in *OsFTL12* rice mutants [50], suggesting an underlying association between OsFLR1/2 and OsFTL12.

Furthermore, the ABA content in *fer-4* mutants is significantly higher than that in wild-type Arabidopsis plants under both control and salt stress conditions [33]. For Si nutrition, its addition may inhibit ABA biosynthesis by down-regulating key ABA biosynthesis-related genes, including *zeaxanthin epoxidase* (*ZEP*) and *9-cis-epoxycarotenoid dioxygenases* (*NCED1* and *NCED4*), when rice plants are exposed to salt stress for a long time (> 24 h) [23]. Given that NaCl-induced ABA negatively regulates the expression of Si transporters [35], ABA levels were measured under various conditions in this study (Fig. 6). Our results demonstrate that both Si and FER are requirement for ABA homeostasis, thereby promoting the growth recovery of salt-stressed rice plants. Moreover, the mitigation of NaCl-induced ABA activity by Si depends on the activation of FER kinases (Fig. 6B), which supports the involvement of OsFLR1/2 in initiating Si signaling, regulating the expression of Si transporters, and facilitating the uptake and accumulation of Si.

As early as 2020, we were among the first to propose that Si alternation of the cell wall — encompassing changes in its composition and mechanical properties — coordinates with cell wall integrity signaling to dynamically regulate plant growth and development in response to environmental changes [14]. This proposition highlighted the potential role of cell wall integrity receptors in Si sensing. Subsequently, several colleagues concurred with our hypothesis and further suggested that mechanoreception molecules on the cell membrane may function as Si sensing receptors [52]. Prior to this study, there was no direct evidence supporting these hypotheses. Our current findings represent a significant advancement in our fundamental understanding of Si signaling initiation. Future research will likely focus on whether other cell wall integrity receptors, such as WAKs, are involved in Si signaling transduction. Additionally, while we have conducted preliminary studies on the relationship between Si nutrition and OsFLR1/2, further investigation is required to elucidate the underlying mechanisms by which OsFLR1/2 functions in Si sensing.

## 5. Conclusions

Our study demonstrates that Si functions as an enhancer of cell wall integrity by regulating pectin methyl-esterification and cellulose microfibril arrangement under salt stress conditions. This regulation modulates the FER homologs OsFLR1/2-mediated cell wall integrity signaling pathway (Fig. 7). Furthermore, Si plays a dual role in modulating the activity of OsFLR1/2 under different stage during salt stress response. In turn, OsFLR1/2 regulates the expression of Si transporter genes, thereby influencing the subsequent uptake and accumulation of Si in salt-stressed rice plants. These findings suggest the existence of a feedback loop between Si nutrition and FER kinase activity, implying the involvement of FLR1/2 in initiating Si signaling.

**Fig. 7.**
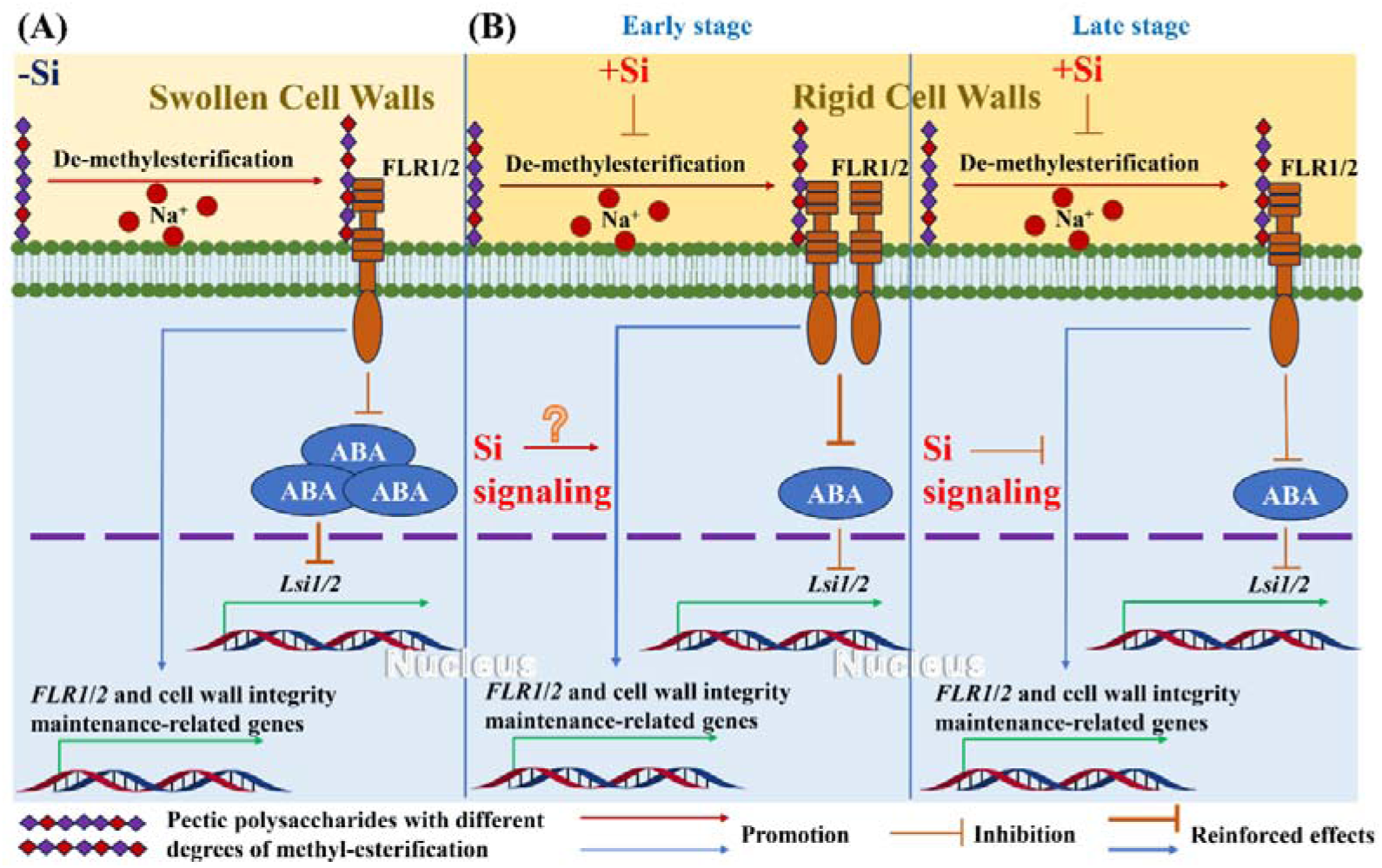
Si mitigates salt stress by modulating pectin methyl-esterification and FLR1/2-mediated cell wall integrity signaling. (A) Rice adaption to salt stress without Si addition. Salt stress induces pectin de-methyl-esterification, leading to cell wall softening and activation of OsFLR1/2-mediated cell wall integrity signaling. Additionally, high concentrations of NaCl increase ABA production, which subsequently inhibits the expression of Si transporter genes *OsLsi1* and *OsLsi2*. (B) The switch roles of Si in regulating the activity of FER homologs OsFLR1/2 in response to salt stress. In the early stage of salt stress, Si application increases pectin methyl-esterification and stiffens the cell wall. Concurrently, Si enhances the activity of OsFLR1/2, although the underlying mechanism remains to be elucidated. In turn, OsFLR1/2 positively regulate the expression of Si transporter genes by inhibiting the synthesis of ABA, thereby promoting Si accumulation. However, in the late stages, Si can suppress the expression of *OsFLR1/2* genes, facilitating growth recovery. The presence of a feedback loop between Si nutrition and FER kinase activity suggests the involvement of OsFLR1/2 in initiating Si signaling.

## Acknowledgements

This work was supported by the Key Research and Development Program of Sichuan Province (Grant No. 2024YFFK0190), the first batch of Scientific and Technological Innovation Team for Qinghai-Tibetan Plateau Research in Southwest Minzu University (Grant No. 2024CXTD04) and the Southwest Minzu University Research Startup Funds (Grant No. RQD2023019).

## Supporting Information

Additional Supporting Information may be found online in the Supporting Information section at the end of the article.

Table S1 Some FT-IR wavenumbers from the main plant cell wall components.

Table S2 Primers used in this study.

