## Supplemental Table 1 and 2 for "Silicon regulation of pectin methyl-esterification and cellulose microfibril organization modulates FERONIA-mediated cell wall integrity signaling for rice adaptation to salt stress" (from another version of this preprint: version 1, 2025-03-19)

**Supporting Information**

**Table S1** Some FT-IR wavenumbers from the main plant cell wall components.

| Assigned cell wall component | Wavenumber (cm^−1^) |
| --- | --- |
| Cellulose | 1050 |
| Hemicelluloses | 1161, 1201, 1313, 1371, 1429, 1442 |
| Pectins | 950, 990 |
| Esterified uronic acid | 1712, 1735 |

**Table S2** Primers used in this study.

| Primer name | Primer sequence (5'-3') |
| --- | --- |
| *OsFLR1*-1-F | GTTGATCCCCACCTGAAGGG |
| *OsFLR1*-1-R | TGACATCCCACATCCAAGGC |
| *OsFLR2*-1-F | GCGCTTGGAGATCTGCATTG |
| *OsFLR2*-1-R | TCGATGGCCCAGTCTTTGAC |
| *OsLsi1*-F | GCCAGCAACAACTCGAGAACAA |
| *OsLsi1*-R | CATGGTAGGCATGGTGCCGT |
| *OsLsi2*-F | ATCACCTTCCCCAAGTTCCT |
| *OsLsi2*-R | CAGCTCCCTCCAGTACATGC |
| *OsHistoneH3*-F | GGTCAACTTGTTGATTCCCCTCT |
| *OsHistoneH3*-R | AACCGCAAAATCCAAAGAACG |
